# Differences in movement and dynamic resource selection between breeders and floaters revealed with an Ornstein-Uhlenbeck space use model

**DOI:** 10.1101/2020.06.15.153429

**Authors:** Joseph M. Eisaguirre, Travis L. Booms, Christopher P. Barger, Stephen B. Lewis, Greg A. Breed

## Abstract

In populations of many taxa, a large fraction of sexually mature individuals do not breed but are attempting to enter the breeding population. Such individuals, often referred to as “floaters” play critical roles in determining dynamics and stability of these populations. Floaters are difficult to study, however, so we lack data on the roles they play in population ecology and conservation status of many species. Here, we paired satellite telemetry and a new mechanistic space use model based on an Ornstein-Uhlenbeck process to study the differential habitat selection and space use of floater and territorial golden eagles *Aquila chrysaetos*. Our sample consisted of 49 individuals tracked over complete breeding seasons across four years, totalling 104 eagle breeding seasons. Modeling these data with the new mechanistic approach was required to parse key differences in movement and separate aspects of resource selection from central place behavior. We found that floaters generally had more expansive space use patterns and larger home ranges, partitioning space with territorial individuals seemingly on fine scales through differential habitat and resource selection. Floater and territorial eagle home ranges overlapped markedly, suggesting floaters use the interstices between territories. Further, floater and territorial eagles differed in how they selected for uplift variables, key components of soaring birds’ energy landscape, with territorial eagles apparently better able to find and use thermal uplift. We also found relatively low individual heterogeneity in resource selection, especially among territorial individuals, suggesting a narrow realized niche for breeding individuals. This work furthers our understanding of floaters’ potential roles in population ecology of territorial species, as well as suggests that conserving landscapes occupied by territorial eagles also protects floaters.

## Introduction

Many long-lived species have extended pre-breeding stages with considerable variation in the age at first reproduction (Zack and Stutchbury, 1992; Gill, 2007). These characteristics often promote a demographic class that is reproductively capable, but members of which do not attempt to breed. In birds, individuals in this demographic group are often referred to as “floaters.” The social dynamics of this class can be highly complex and ordered but receives little attention from biologists who have tended to focus on breeding individuals (Penteriani et al., 2011). The idea of a structured but unseen “underworld” within the floater demographic class was introduced as a concise metaphor for their behavioral dynamics and interactions (Brown, 1969; Smith, 1978).

Members of this underworld are individuals that do not breed because of saturated breeding territories and/or fitness trade-offs between occupying low versus high quality territories (Brown, 1969; Newton, 1992; Zack and Stutchbury, 1992; Ens et al., 1995; Kokko and Sutherland, 1998; Newton, 1998; Hunt, 1998; Penteriani et al., 2003; Ferrer et al., 2015). Floaters were initially hypothesized as nomadic individuals that passively await an available territory or poor competitors excluded from territories (i.e. a “doomed surplus”; Krebs, 1971; Stutchbury and Robertson, 1985; Eckert and Weatherhead, 1987; Newton, 1992). However, floaters have been shown to be capable competitors simply employing a different strategy (Smith, 1978; Smith and Arcese, 1989), and their presence and behavior are often important to population dynamics and evolutionary processes (Kokko and Sutherland, 1998; Hunt, 1998; Penteriani et al., 2005, 2011; Lee et al., 2017). The stability of some populations may even be more sensitive to changes in population vital rates of floaters than breeders (Hunt, 1998; Penteriani et al., 2011), and discounting floaters in demographic studies can mask low population growth rates (Penteriani et al., 2011; Lee et al., 2017; Hunt et al., 2017).

Populations of slow-maturing species that lack floaters, or that simply lack floaters adjacent to territories, may be more susceptible to variation in mortality of reproductive adults. Younger and less mature individuals could replace losses of breeding-age individuals, but these less mature, inexperienced individuals would be of lower quality and many would be incapable of actually breeding successfully (Carrete et al., 2006b). This situation would decrease overall reproductive performance at the population level, magnifying impacts of breeder mortality. Floaters in a population can buffer against such elevated breeder mortality, as they are experienced, can rapidly fill territory vacancies, and thus maintain reproductive success of the population (Hunt, 1998; Penteriani et al., 2011). Although many floaters have not yet bred for the first time, all have spent at least one, but in some cases many, years as floaters before recruiting to the breeding class, which affords experience in foraging and survival as well as more time to physically mature. Depending on how floaters use space, they may have persisted within or near breeding territories for some time and accrued experience with its resources and risks that will increases the odds of success once an opening becomes available (Stamps, 1995; Fagan et al., 2013). Such information can improve realized fitness (Forrester et al., 2015), leaving them primed to immediately perform as capable breeders and act as a demographic buffer. Further, within the underworld social structure, experience also likely helps floaters compete against other floaters, with better competitors taking over territories when they become available.

There are essentially three basic space use strategies employed by floaters (Newton, 1976, 1979; Penteriani et al., 2011). They may (1) geographically segregate from breeders, (2) establish along the fringes of breeding habitat, or (3) persist in the interstices among and occasionally within breeding territories. A floater utilizing strategy 3 would have access to information and experience with candidate territories, but would also have to cope with conflict from defending territorial individuals. Individuals employing strategy 1, by contrast, would establish a presence near to, but segregated from, viable territories while awaiting a territorial opening, thereby minimizing agonistic interactions with territorial individuals (Hunt, 1998; Caro et al., 2011; Penteriani et al., 2011). However, floaters may still prefer habitat and/or have requirements otherwise similar to breeders, but lacking a central place to defend. Alternatively, they could also have different habitat preferences. In either case, floaters and breeders would be spatially segregated, reducing the territorial defense requirements of breeders but leaving floaters with less information and experience about the resources available in territories. Further, if left undiscovered, spatial segregation between floaters and breeders could lead to ineffective conservation or management actions, as any action would likely be focused on habitats used by breeding individuals (Penteriani et al., 2005, 2011).

When floaters persist along the fringes of breeding habitat or in the interstices of breeding territories—strategies 2 and 3—they gain knowledge of habitat and territories but may affect territorial individuals and put themselves at risk. More frequent territory defense against floaters could potentially reduce reproductive success (López-Sepulcre and Kokko, 2005; Carrete et al., 2006a; Bretagnolle et al., 2008). Floaters may also affect the survival of territorial individuals through fatal conflict; some have hypothesized that survival of territory-holders may be lower than floaters’ due to repeated territorial conflict elicited by floaters (Hunt, 1998). We would expect these effects on breeding behavior, success, and survival to be greatest when floaters persist in the interstices of territories, likely making occasional intrusions into territories as well (strategy 3). However, this strategy would simultaneously allow floaters to gain the most knowledge of candidate territories. In contrast, floaters persisting on the fringes of breeding habitat (strategy 2), possibly excluded by breeders, would relax defense requirements of breeders, but would allow comparatively limited opportunity for floaters to learn about territories.

Spatial learning in home ranges and territories could benefit certain taxa more than others (Fagan et al., 2013). For example, soaring birds rely on dynamic energy landscapes (Shepard et al., 2013) in the form of local upward air currents (i.e. uplift) to offset the energetic expense of flight, and these flight subsidies can drive soaring bird movement and behavioral processes (Shepard et al., 2011, 2013; Katzner et al., 2015; Miller et al., 2016; Eisaguirre et al., 2018, 2019a). Given that topography and atmospheric process vary spatially on fine scales, the energy landscape, in addition to other key resources such as habitat and food availability, likely varies among territories. Thus, learning territory-specific energy landscapes could allow individuals to move around a territory more efficiently and quickly (Stamps, 1995), improving resource acquisition and risk avoidance, as well as increasing the probability of reproductive success and thus realized fitness. We might therefore expect that floater space use patterns and overall floater strategy in soaring birds and other taxa could differ from breeders with respect to use of dynamic energy landscapes.

How floaters use space and their social dynamic within a population likely plays a key role in how they impact population dynamics and their ability to act as a buffer against breeder mortality (Hunt, 1998; Penteriani et al., 2011; López-Sepulcre and Kokko, 2005; Carrete et al., 2006a; Bretagnolle et al., 2008). Moreover, although often overlooked, understanding these individuals may be considerably more important in assessing conservation status and developing conservation strategies than is often assumed (Penteriani et al., 2011). However, few detailed investigations into the spatial ecology of floaters have been undertaken (Rohner, 1997; Campioni et al., 2012; Tanferna et al., 2013; Penteriani et al., 2015). This is due in part to the difficulty of studying this segment of the population, as floaters are often secretive as to avoid conflicts with breeders and, unlike breeders, lack spatial fidelity to specific locations on the landscape (i.e. nests; Sergio et al., 2009; Penteriani et al., 2011).

Technological advances in animal telemetry now permit collection of detailed data throughout substantial periods of animals’ lives. Such data can be paired with appropriate statistical methods recently developed for inferring animal behavior from telemetry data (many reviewed by Hooten et al., 2017). With such tools available, we are now better positioned to understand the secretive underworld of floaters, how they interact with their territory-holding conspecifics, and infer the potential importance of this demographic class to population stability and dynamics.

Ornstein-Uhlenbeck (OU) processes have long been used to model the movement and space use of animals and may be useful for studying the comparitve movement ecology of floaters and breeders (Dunn and Gipson, 1977; Blackwell, 1997; Johnson et al., 2008; Breed et al., 2017). The OU process naturally captures movement around a central place and, when coupled with a resource weighting function, can account for and allow inference about an animal’s preferences for features on heterogeneous landscapes, such as habitats and dynamic uplift (Johnson et al., 2008; Eisaguirre et al., 2020). Modelling approaches based on the OU process have been developed to allow for multiple home range or territory core areas, which might be expected for an animal with one or multiple established foraging areas, in addition to a nest or den (Johnson et al., 2008; Breed et al., 2017; Eisaguirre et al., 2020). Appropriately implemented, such models could be quite useful for revealing how floater and breeder movement and space use strategies vary based on central places as well as static and dynamic landscape features.

Here, we used a large data set on the movements of individual breeding-age golden eagles *Aquila chrysaetos* collected with GPS telemetry in southcentral Alaska, coupled with a newly developed mechanistic OU space use model, to determine how floaters use space, resources, and dynamic energy landscapes as compared to breeding individuals. Specifically, we investigated whether floaters (1) are geographically segregated from territorial breeders, (2) establish along the fringes of breeding habitat, or (3) persist in the interstices of established and occupied breeding territories. Golden eagles are a long-lived, territorial species, populations of which can consist of large segments of floaters about which little is known (Watson, 2010), making them an excellent study system for understanding the behavior and potential importance of this poorly understood demographic class to population dynamics across large spatial scales. Further, as golden eagles are soaring birds, we additionally accounted for and investigated how energy landscapes influenced floater and breeder space use dynamics.

## Methods

### Study area & system

Golden eagles are widely distributed, exhibit high territory fidelity, and most commonly nest on cliffs in many parts of their holarctic range (Kochert et al., 2002; Watson, 2010). Golden eagles are also generally central place foragers (Kochert et al., 2002; Watson, 2010), so suitable nesting habitat must lie within or adjacent to suitable foraging habitat. The number of viable territories is, therefore, limited in many areas, which has given rise to sometimes large floater sectors of populations, about which we know very little (Watson, 2010).

In many northern populations, including our study population, golden eagles behave as long distance migrants (Watson, 2010). Eagles in our sample summered primarily in the western Alaska Range, Talkeetna and Chugach Mountains of Alaska, USA and overwintered in the Rocky Mountain West, including Colorado, Utah, Wyoming, Montana, Idaho, Oregon, and Washington, in the US and mid to southern Alberta and British Columbia in Canada (Eisaguirre et al., 2019a). The summering area is a diverse matrix of habitats, including boreal forest, open tundra, high alpine, and glaciers. Aerial surveys of the nest sites of tagged eagles indicated that all nested on cliffs, except for one that nested in a tree.

### Telemetry data

We captured golden eagles with a remote-fired net launcher placed over carrion bait near Gunsight Mountain, Alaska (61.67°N 147.35°W). Captures occurred during spring migration, 15 March to 15 April 2014-2016. Our capture period overlapped the time when territory holders would be returning to their territories. Adult and sub-adult eagles were equipped with 45-g back pack solar-powered Argos/GPS platform transmitter terminals (PTTs; Microwave Telemetry, Inc., Columbia, MD, USA). Eagles were sexed molecularly and aged by plumage.

PTTs were programmed to record GPS locations on duty cycles, ranging from 8-14 fixes per day during the breeding season, depending on year of deployment. In 2014, PTTs were set to record 13 locations at one-hour intervals centered around solar noon plus a location at midnight local time. 2015 PTTs were programmed to record eight locations with one-hour intervals centered around solar noon very early and late in the season and 10 locations for most of the season. In 2016, we revised our programming approach so that PTTs took eight fixes daily early and late and 12 fixes most of the season with a fixed 3-/2-hr time interval. Fifteen PTTs were deployed in 2014, 23 in 2015, and 15 in 2016.

### Mechanistic space use model

Given that golden eagles are generally central place foragers and rely on dynamic energy landscapes (Watson, 2010), we adapted and implemented a recently introduced approach designed for making practical, mechanistic inference about animal space use while considering such patterns (Eisaguirre et al., 2020). This approach uses a model with core dynamics based on the Ornstein-Uhlenbeck (OU) process (Dunn and Gipson, 1977; Blackwell, 1997). Although a continuous-time process, the inherently discrete nature of data collected with GPS telemetry makes the position likelihood of the OU process, which takes the form of an OU biased random walk (BRW), most useful:

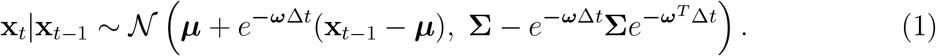

Here, **x**_*t*_ is a coordinate vector of the location of the animal at time *t*, ***ω*** = *ω***I**_2_ with *ω* describing the strength of the animal’s tendency toward the animal’s central point ***μ***, **Σ** = *σ*^2^**I**_2_, and *σ* > 0. The particular isotropic form presented here, with a single centralizing parameter *ω* and diffusion parameter *σ*, gives rise to the steady-state space use distribution 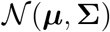, contours of which are circular about ***μ***. The OU parameters naturally lend themselves to direct biological interpretation. As *σ* is the diffusion parameter of the OU process and *ω* is the intensity of the attraction to the home range center, larger values of *σ* and smaller values of *ω* equate to larger, more diffuse home ranges. In contrast, smaller values of *σ* and larger values of *ω* equate to smaller, more concentrated home range cores.

The distribution 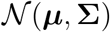 is what a bird’s space use distribution would look like while tending a nest surrounded by uniformly distributed average habitat. Nonuniform and possibly dynamic resources plus a bird’s preferences for those resources would give rise to a time-varying modified form of that distribution though. This can be accounted for with a resource weighting function, such that we can write the conditional probability density of the animal’s location at time *t* (Johnson et al., 2008):

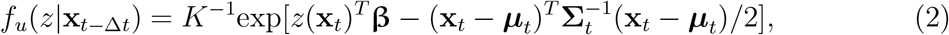

where *z* is the landscape over which the animal is moving, *z*(**x**_*t*_) corresponds to a vector of environmental (e.g., weather and habitat) variables at **x**_*t*_ and time *t*, **β** weights the elements of *z*(**x**_*t*_) based on the animal’s preferences, ***μ***_*t*_ = ***μ*** + *e*^−*ω*Δ*t*^(**x**_*t*−Δ*t*_ − ***μ***), and 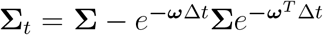. The steady state form of equation 2 is simply the product of an exponential weighting function and 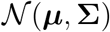 (Eisaguirre et al., 2020).

As an eagle’s space use could be centered around multiple locations, rather than just a single nest, such as roosts or hunting locations (Watson, 2010), we accounted for the presence of multiple central points in estimating eagle space use, following Eisaguirre et al. (2020). Given we did not have fixed time intervals for all tags deployed on eagles, we forewent modeling transitions as a Markov process (*sensu* Eisaguirre et al. 2020), and simply used an indexing approach that did not estimate transition probabilities among *K* many cores (*sensu* Johnson et al., 2008). Note that utilizing the OU model weights the use of central points, such as nest sites and perches, independent of habitat selection, which serves to reduce any overestimation of the importance of habitat or terrain features possibly correlated with the locations of those central points.

### Weather and habitat variables

#### Static landscape variables

We used the Alaska Center for Conservation Science Alaska Vegetation and Wetland Composite (AKVWC; 30 m resolution) data for characterizing habitat type. We collapsed the numerous habitat types in the data set to eight for this analysis. These were shrub, open (but vegetated; e.g., meadows and open tundra), bare, forest, wet (e.g., marsh), water, ice (i.e. perennial snow and ice), and human. See Appendix 1 for details.

Elevation data were gathered using the Mapzen Terrain Service with the `elevatr` package (Hollister and Shah, 2018). We specified the ‘zoom’ variable such that the resolution closely matched that of the habitat data. We included elevation and slope (*slope* ∈ [0, *π*/2] radians) as predictors in the model.

#### Dynamic landscape variables

We used a state-wide data set of snow-off date (date of which an area became snow free) to derive a dynamic binary indicator variable of whether or not grid cells were free of snow (Macander et al., 2015). While one might expect some confounding between the (perennial) snow and ice habitat variable and this snow indicator, it would be limited due to few glaciated and perennial snow-covered areas frequented by eagles sampled.

The remaining variables included in the model were related to orographic and thermal uplift and were derived from the elevation data and Center for Environmental Predictions (NCEP) North American Regional Reanalysis (NARR) data. Angle of incidence was included for the effect of orographic uplift on eagle space use. It is the deviation of the relative wind from the aspect of a slope and was computed such that *aoi* ∈ [0, *π*] (Murgatroyd et al., 2018); *π/*2 corresponds to a wind orthogonal to a slope, and *π* a wind perfectly parallel blowing up slope. Wind direction was computed trigonometrically from the meridional and zonal wind components estimated by the NCEP NARR 10 m above the surface.

The effect of thermal uplift was included with a hill shade variable. Hill shade was computed following Murgatroyd et al. (2018), such that *hs* ∈ [0, 1], and we gathered the required location-, date-, and time-specific azimuth and zenith of the sun using the package maptools (Bivand and Lewin-Koh, 2016). Higher hill shade corresponds to more direct sun and thus more potential for thermal uplift.

### Inference

We fit the mechanistic space use model to four breeding seasons of data, 2014-2017, with Stan and R following Eisaguirre et al. (2020) (Stan Development Team, 2016, 2018; R Core Team, 2018). We estimated the movement and selection parameters hierarchically across individuals (details presented by Eisaguirre et al. 2020), but for computational reasons we fit the model separately for certain cases.

When the nestling(s) of a breeding pair mature enough to thermoregulate (about three weeks after hatching), the female, who does the majority of incubating and parental care, is no longer required to care for them as regularly, so she is free to move more widely about the territory and help with provisioning (Watson, 2010). We hypothesized that when this occurs, space use may change substantially due to members of a pair partitioning space. We expected this might also impact floater movement as well due to the female being more free to defend the territory against floating intruders. Thus, within each year, we fit the model separately for the four combinations of floater or territorial and early or late breeding season. Aerial observations of the nests of tagged individuals indicated that 20 June was, on average, the date nestlings were about three weeks old, so we used this date to partition the data into early and late periods of the breeding season.

Distinguishing floater from territorial eagles was done a priori by visual inspection of each individual’s movement data. As we suspected eagles might use resources and habitat differently in different home range cores, we also estimated core-specific effects for each predictor. All continuous predictors were centered and standardized to mean zero and unit variance for estimation. Additionally, given that eagles had different numbers of home range cores and core-specific movement parameters, we chose to summarize and interpret the OU movement parameter estimates for the most used home range core (i.e. that in which an eagle spent the most time) for each eagle breeding season to help understand movement and home range behavior (Dunn and Gipson, 1977).

Lastly, given the properties of the OU model, we were able to compute analytical estimates of eagle space use distributions (Eisaguirre et al., 2020). We used a Bayesian hierarchical Gamma regression (2000 iterations, default priors; Stan Development Team, 2016) to assess the differences in home range size between floaters and territorial eagles and early and late breeding season. To be comparable with other home range estimation methods, we defined our home range estimator as the 95% contour of the utilization distribution (Hooten et al., 2017). Year and individual were included as random effects to account for repeated measures. Differences were assessed by constructing each respective posterior predictive distribution.

## Results

We were able to determine both arrival and departure dates in at least one year for 49 individual eagles between 2014-2017 (Table 1), and thus our total data set consisted of 104 eagle breeding seasons. Of those, 78 were territorial (44 male and 34 female) and 26 were floaters (24 male and 2 female).

**Table 1:** Median (interquartile range in days) dates of arrival on and departure from the summer range of golden eagles tagged with GPS telemetry in southcentral Alaska.

|  | 2014 | 2015 | 2016 | 2017 |
| --- | --- | --- | --- | --- |
| **Territorial** |  |  |  |  |
| Arrival | 29 March (5) | 27 Mar (4) | 26 Mar (8) | 1 Apr (6) |
| Departure | 28 Sep (28) | 7 Oct (17) | 5 Oct (13) | 4 Oct (18) |
| **Floater** |  |  |  |  |
| Arrival | 5 Apr (3) | 30 Mar (6) | 24 Mar (10) | 1 Apr (2) |
| Departure | 24 Sep (12) | 24 Sep (4) | 29 Sept (4) | 6 Oct (13) |

Sixteen of the floater breeding seasons were from eagles aged as entering their fifth year at capture, six from eagles entering some year after their fourth year, two after their fifth year, and two as ‘adult’ (likely after fifth year). Nearly all territorial eagle breeding seasons were from individuals aged as after their fifth year at capture, but six were fifth year, one after fourth year, and two as ‘adult’ (likely after fifth year).

### Movement parameters

Floaters and territorial eagles established similar numbers of home range cores, but it seemed territorial eagles established slightly more cores later in the breeding season (Fig. 1). OU movement parameters varied markedly between floaters and territory holders (Figs. 1). Compared to floaters, territorial eagles had smaller home range cores, indicated by smaller movement variance 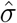 (Fig. 1), and stronger attraction toward the center of their home range core, indicated by larger 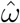 (Fig. 1). Additionally, eagles’ most used cores were smaller during late breeding season (Fig. 1).

**Figure 1:**
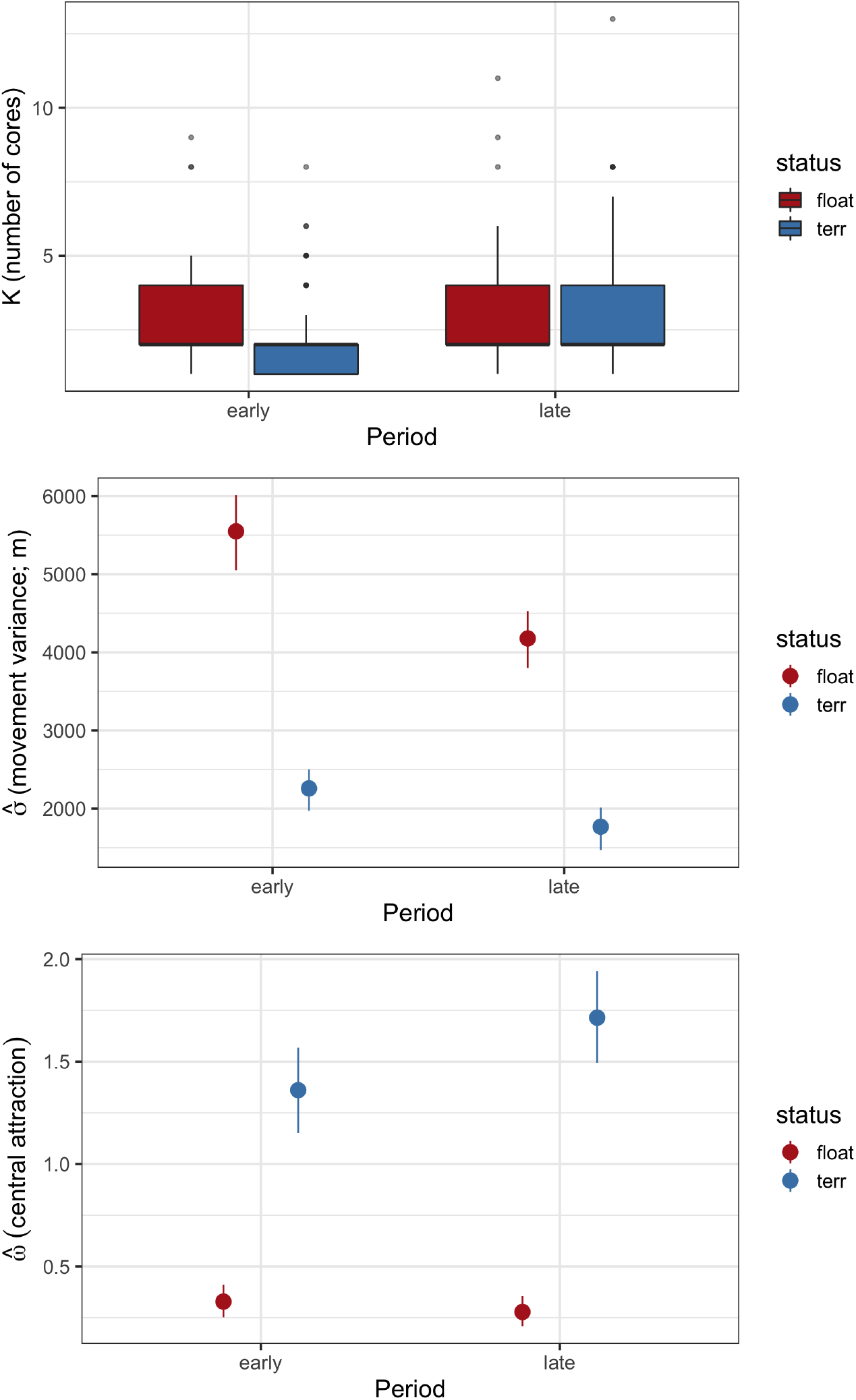
Summaries of the number of home range cores *K* and population-level estimates (posterior means and 90% credible intervals) of the movement variance parameter *σ* and centralizing tendency parameter *ω* from the Ornstein-Uhlenbeck space use model for golden eagles summering in southcentral Alaska. OU parameters are for the most used home range core, and the model was fit separately for early and late breeding.

### Habitat selection

#### Effects of static landscape variables

Both floater and territorial eagles selected for higher elevations and steeper slopes (Fig. 2). Four of the habitat types (bare, open, forest, and shrub) comprised over 99% of the habitats used by both territorial and floater eagles, so we report and interpret the results for those here. The habitat types territorial and floater eagles selected were generally similar (Fig. 2 & 3), but there were a few important differences. Floaters used forested areas more than territorial eagles and much less during late breeding season than early (Fig. 3). Additionally, floaters more strongly selected for shrub and open areas than territorial eagles, and territorial eagles used bare areas more frequently than floaters (Fig. 2 & 3).

**Figure 2:**
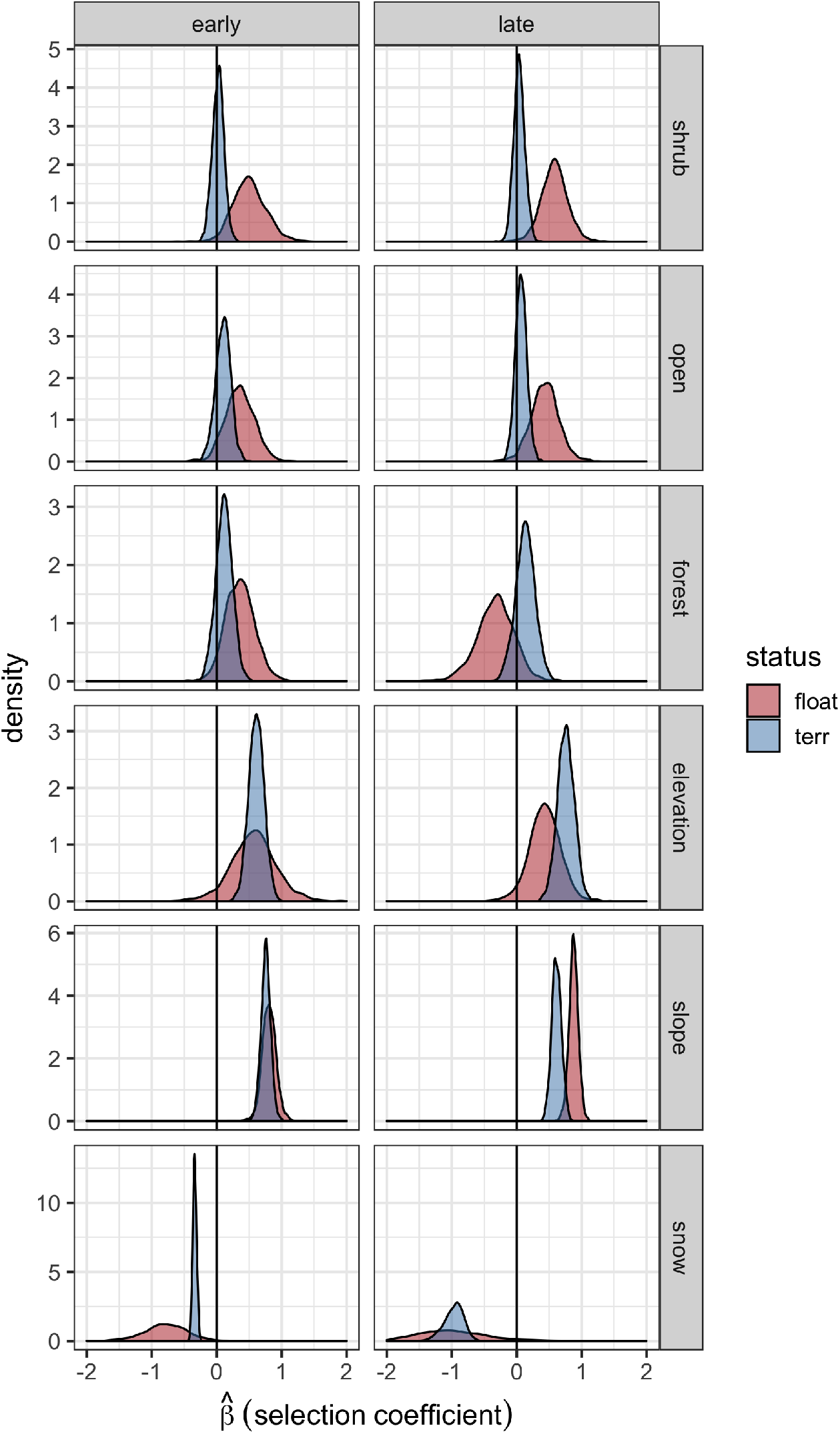
Estimated marginal posteriors for the population-level selection coefficients from an Ornstein-Uhlenbeck space use model fit to summer golden eagle GPS data in southcentral Alaska. The model was fit separately for early and late breeding season and floater and territorial eagles. ‘bare’ (bare ground) is the reference category.

**Figure 3:**
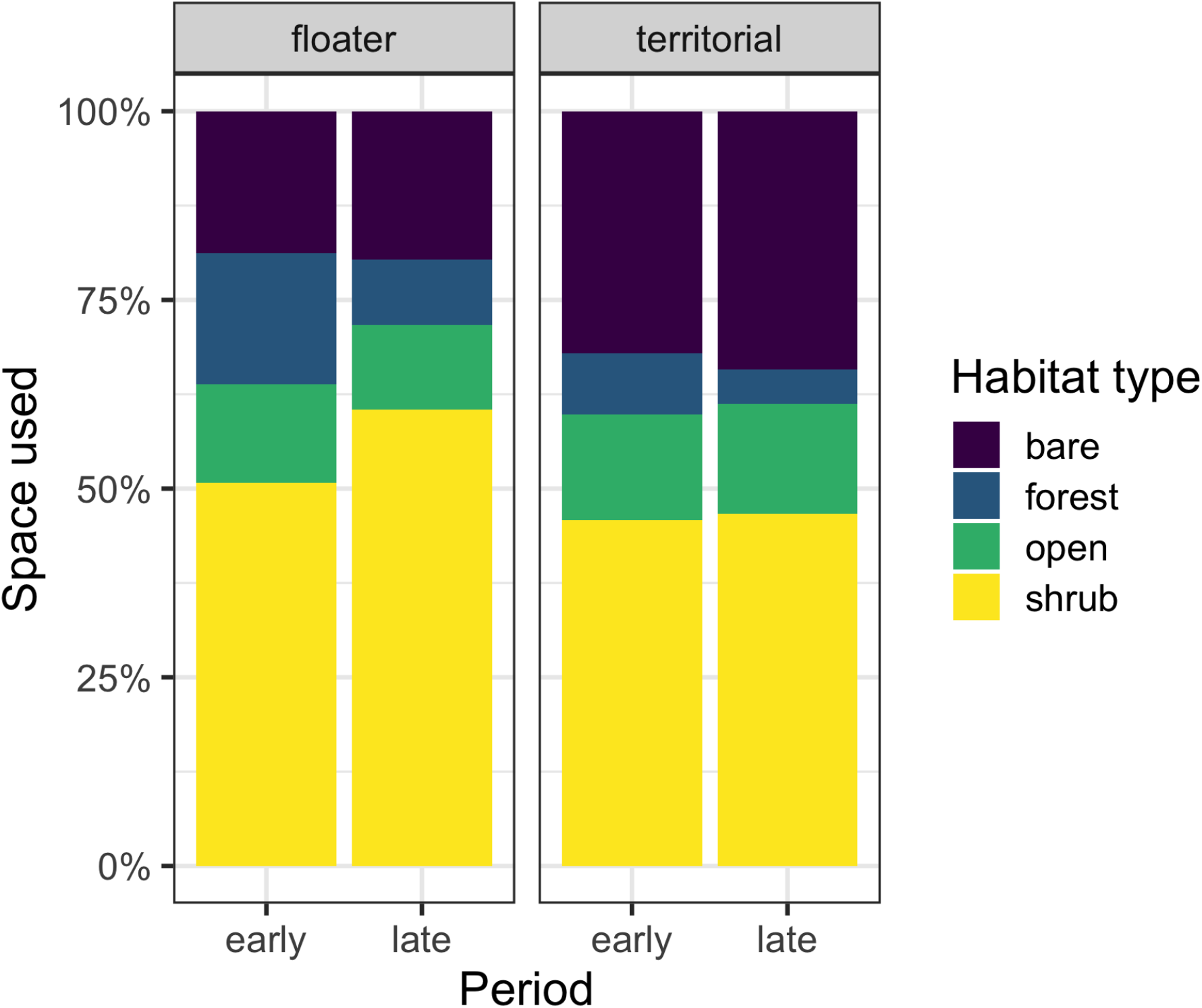
Proportion of habitat types used by golden eagles summering in southcentral Alaska.

#### Effects of dynamic landscape variables

Both territorial and floater eagle space use was affected by thermal and orographic uplift nearly identically (Fig. 4). However, territorial eagles exhibited a stronger preference for areas with more thermal uplift than floaters (Fig. 4). Both territorial and floater eagles selected strongly against snow cover during late breeding season with generally weaker selection against it during early breeding season (Fig. 2).

**Figure 4:**
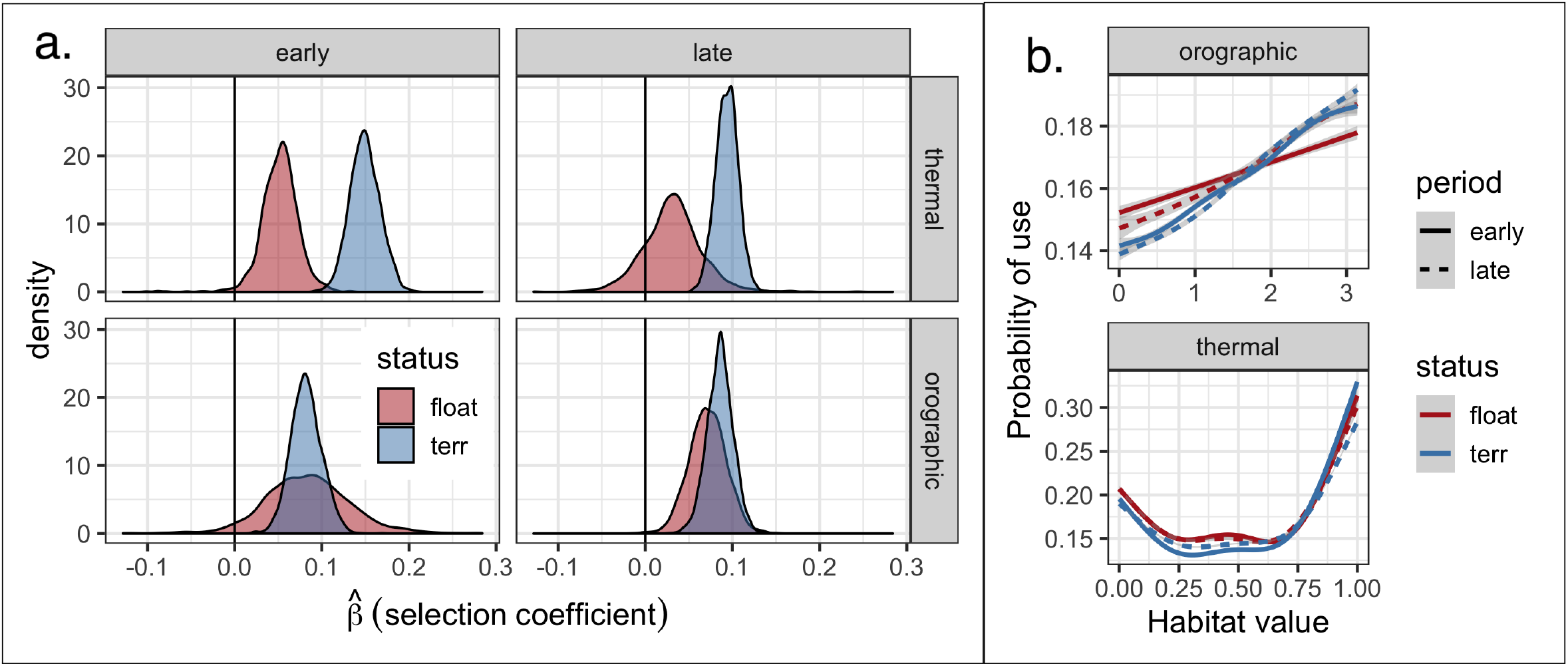
(a.) Estimated marginal posteriors for the population-level selection coefficients for golden eagles in southcentral Alaska estimated with an Ornstein-Uhlenbeck (OU) space use model, and (b.) probability of a golden eagle using a spatial location within its breeding season home range in southcentral Alaska as a function of habitat variables. The model was fit separately for early and late breeding season and floater and territorial eagles. In b. predictions were smoothed over the availability points with a generalized additive model (*df* = 6) and ribbons are 95% confidence intervals. Angle of incidence (measured in radians) was used to proxy orographic uplift, and hill shade was used to proxy thermal uplift. Note that higher hill shade corresponds to more direct sun and thus greater thermal uplift potential.

#### Individual and home range core variance

Unlike in the OU movement parameters, there was relatively low among-individual variance in habitat selection parameters, especially for territorial individuals (Fig. 5). There was, however, more individual heterogeneity in selection parameters for floaters than territory-holders (Fig. 5). In contrast, among-core variability in selection parameters was markedly high (Fig. 5), suggesting eagles exhibited different habitat selection patterns in different home range cores. This difference between individual and core variance was further supported by permutation tests on paired medians of the variance point estimates (*p* << 0.001).

**Figure 5:**
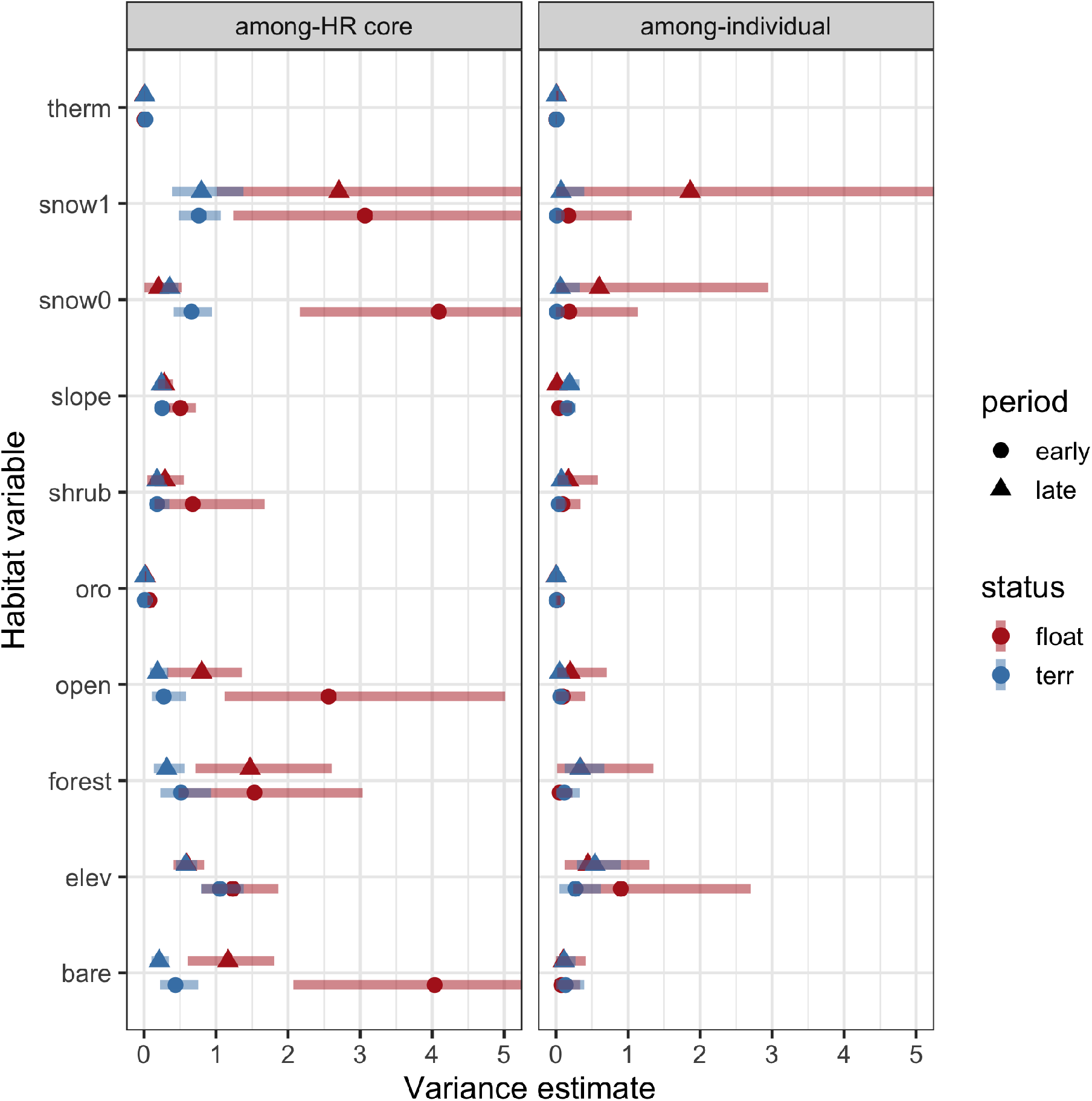
Estimated among-home range core variance and among-individual variance for the selection coefficients from an Ornstein-Uhlenbeck space use model fit to summer golden eagle GPS data in southcentral Alaska. The model was fit separately for early and late breeding season and floater and territorial eagles. Points are posterior means, and horizontal lines are 95% credible intervals. Note the horizontal axis has been truncated for presentation.

Further, both the estimated among-individual and among-core variances for thermal and orographic uplift were very low (Fig. 5). This suggests strong similarity in how different eagles respond to these variables as well as similarity in how those eagles respond within different home range cores.

### Realized home range size

Due to the properties of the OU model, estimating home ranges with the model can be done by simply taking the product of the OU and habitat weighting function steady state distributions (Eisaguirre et al., 2020). Applying this, here, we found that floaters had larger home ranges than territorial eagles during both early and late breeding season (Fig. 6). Additionally, both territorial and floater eagles exhibited larger home ranges early in the breeding season compared to late (Fig. 6), consistent with the pattern in home range core size mentioned above (Fig. 1). Median (IQR) home range sizes for territorial eagles were 31 km^2^ (18, 143) and 24 km^2^ (13, 53) for early and late breeding season, respectively, and 581 km^2^ (326, 704) and 346 km^2^ (143, 496) for floaters.

**Figure 6:**
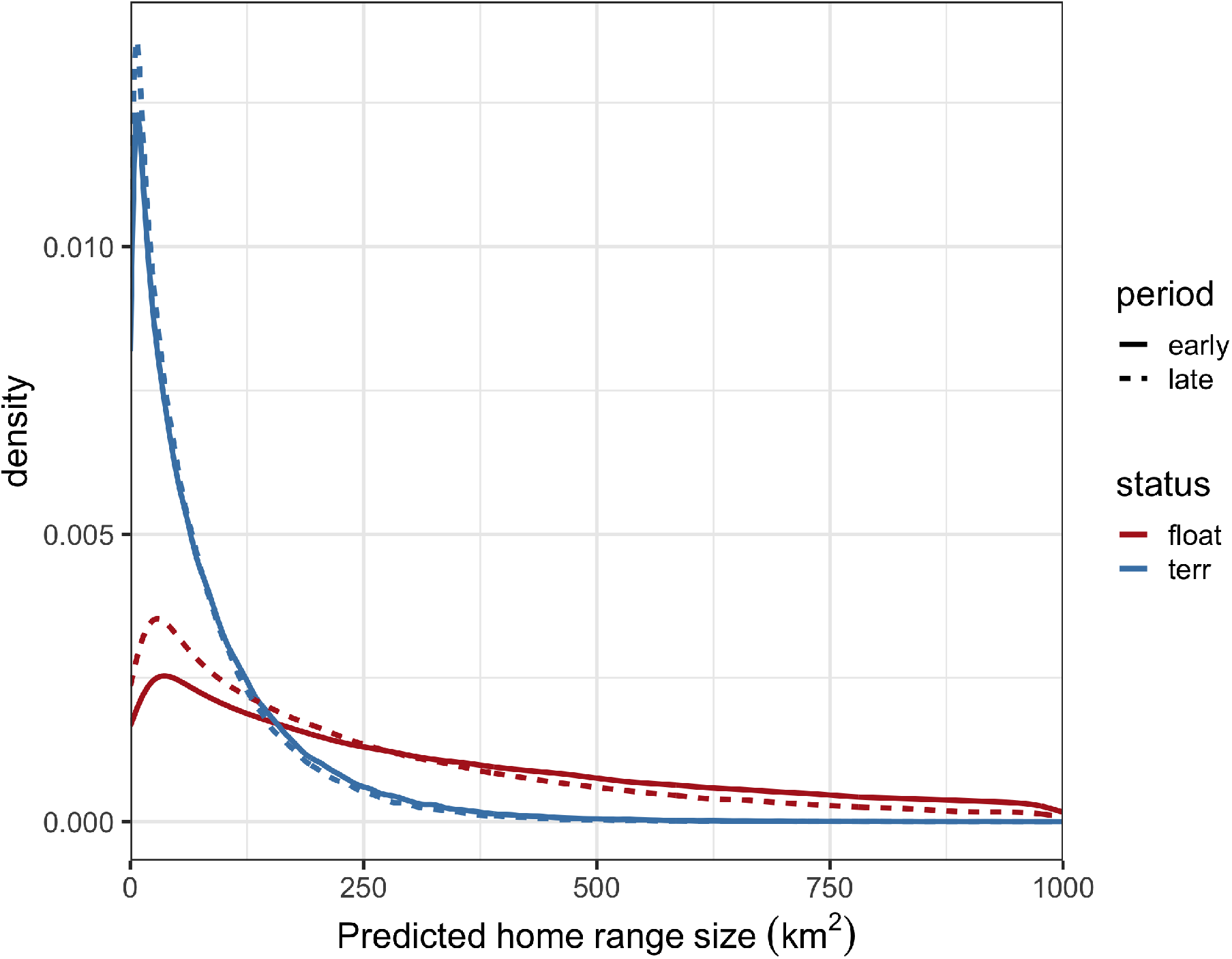
Posterior predictive densities of a hierarchical Gamma regression on golden eagle home range size predicted with an Ornstein-Uhlenbeck home range model for golden eagles summering in southcentral Alaska 2014-2017. Period of breeding season (early or late) and status (floater or territorial) were included as fixed effects, and individual and year were included as random effects.

## Discussion

Here we implemented a newly developed mechanistic movement model to investigate differences in space use between territorial and non-territorial floater individuals. This approach allowed us to parse the relative importance of three different landscape features on movement within home ranges (affinity to a central place, static landscape features, and dynamic energy subsidies) to understand how each affected space use of territorial breeding individuals and floaters. The model allowed us to investigate differences in movement behavior, habitat and dynamic resource selection, and home range size while considering that individuals can structure their home ranges with multiple core areas as well as move and select resources differently in different cores.

We found that floater and territorial golden eagles exhibited slight differential selection for habitat and resources, but floater space use patterns were much more expansive, consistent with floater behavior in other taxa (e.g., great horned owl *Bubo virginianus* and black kite *Milvus migrans*; Rohner, 1997; Tanferna et al., 2013). Although we sampled a relatively small proportion of golden eagles that summer in southcentral Alaska, an expanded view of some neighboring tagged individuals (Fig. 7) suggests many floaters use much of the same or closely adjacent space to territorial eagles. The data broadly supports the existence of a demographic underworld of floater eagles (*sensu* Smith, 1978), likely employing strategy 3 and persisting in the interstitial spaces among and within breeding territories.

**Figure 7:**
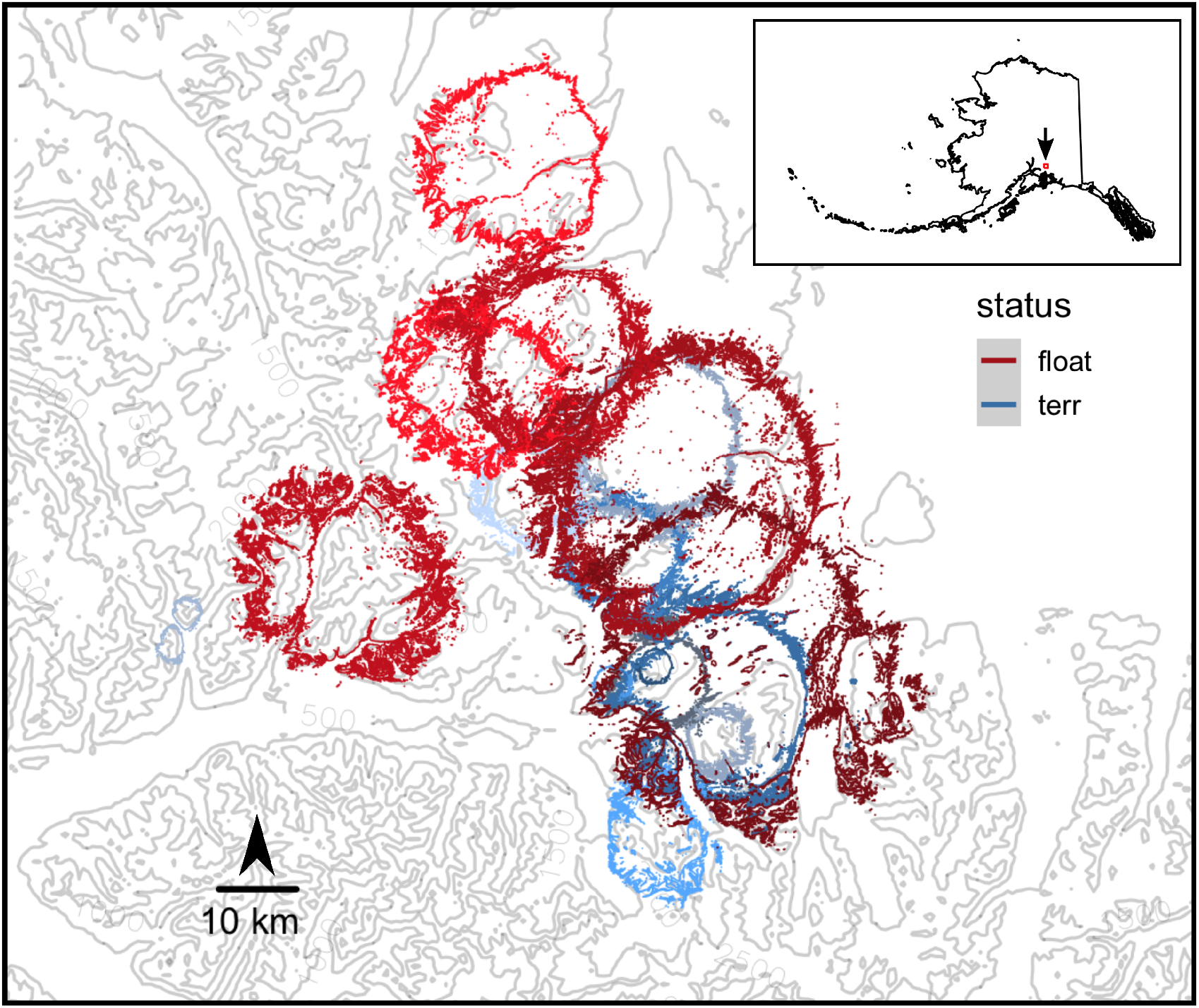
Estimated home range boundaries for nine territorial (blue hues) and four floater (red hues) golden eagles in southcentral Alaska during late breeding season in 2016 plotted over elevation contours. Home range estimates are the 95% contour of the utilization distribution constructed from the posterior predictive distribution of an Ornstein-Uhlenbeck space use model.

For very long-lived species, a strategy of persisting amongst breeding territories may be particularly adaptive. Gaining knowledge of habitat and resource distributions within territories prior to occupation and breeding attempts would improve individuals’ initial reproductive success (Stamps, 1995). In highly competitive environments, this advantage should outweigh the risks associated with constant contention with breeders and physical interactions likely required for displacing territory-holders. Further, our results suggest the floating period includes learning about not only static habitats and terrain but dynamic features of the energy landscape as well (Fig. 4). That floaters showed weaker selection for thermal uplift than territorial eagles may be attributed to floaters being less familiar with both when and where these dynamic resources are available within and around candidate territories. In contrast, territorial eagles have usually maintained their territory for several breeding seasons and thus have commensurate experience with their territory as well as the nuances of its resource availability.

### Use of energy landscapes within home ranges

The effects of uplift and wind on the behavior and movement of soaring birds is well established (e.g., Shepard et al., 2011; Katzner et al., 2015; Péron et al., 2017; Eisaguirre et al., 2018, 2019a), but less work has focused on how these variables affect emergent individual- and population-level space use patterns (Shepard et al., 2013; Watson et al., 2014; Eisaguirre et al., 2019b). Our finding that golden eagles use space within their home ranges in accordance with thermal uplift contrasts with studies that have used static variables to proxy thermal uplift (Watson et al., 2014). Accounting for the inherently dynamic nature—both across seasons and within days—of thermal uplift, as we did by computing spatio-temporally explicit hill shade, is important to drawing correct inferences about soaring bird space use.

Use of orographic uplift is key for soaring taxa to find food and patrol territories (McLeod et al., 2002; Harmata, 1982; Collopy and Edwards, 1989). The benefits and utility of using orographic uplift is likely not restricted just to territorial individuals, however, as floaters too need to find food and patrol territory boundaries looking for a vacancy or usurping opportunity. We found that orographic uplift drove space use similarly for floaters and territorial eagles (Fig. 4).

### Terrain & habitat use

Topography affects the movement and space use of a range of taxa (Boyce et al., 2003; Shepard et al., 2013), but for eagles, it has been hypothesized that its effect is primarily through interaction with wind (i.e. orographic uplift; McLeod et al., 2002). However, in addition to the dynamic effects of orographic uplift that we found, we also found selection for higher elevations and steeper slopes (Fig. 2), which is consistent with findings for golden eagles elsewhere in North America (Watson et al., 2014). These patterns were also generally similar between territorial and floater eagles, supporting our conclusion that there is not large scale spatial segregation (e.g., along an elevation gradient) between the two groups.

Floaters are more free to select space based on prey abundance due to the lack of nesting and territorial defense requirements (Penteriani et al., 2011). During early breeding season, we found floaters tended to select and use forested areas in addition to shrubbier and open habitats (Fig. 2 & 3). Much of the early breeding season takes place before the emergence of hibernating Arctic ground squirrels *Urocitellus parryii*, which are a primary prey of golden eagles in Alaska (McIntyre and Adams, 1999; McIntyre and Schmidt, 2012; Herzog et al., 2019). During that time, floaters may be favoring forested, shrub, and/or edge habitats slightly, to optimize their access to snowshoe hare *Lepus americanus* and ptarmigan *Lagopus* spp., two other important prey, while still maintaining their proximity to candidate territories.

### Home range structure

Finding that floaters used more space and were less tied to central points than territorial individuals (Fig. 1 & 6) was not surprising, and it is consistent with floaters’ ability to move more freely about the landscape without tending a nest or young. Furthermore, territorial individuals likely do not tolerate floaters within or near territory boundaries for long (Watson, 2010), causing more expansive movements and perhaps ‘pinballing’ among territories. Our results also suggest that the more expansive movements of floaters were influenced by differential habitat use and resource selection (Fig. 2). The higher elevation, barer terrain with suitable nest cliffs is primarily occupied by territorial individuals. Resident pairs’ agonistic behavior toward floaters suppresses their use of these primary areas, leaving floaters to occupy the interstices between territories or, temporarily, areas within territories (i.e. before being expelled by a resident pair; Fig. 7).

Other species of eagles are known to establish multiple home range cores (Watson, 2010), as do a wide range of other other taxa (e.g., sea otter *Enhydra lutris*, black rhinoceros *Diceros bicornis minor*, and land iguana *Conolophus pallidus*; Breed et al., 2017; Lent and Fike, 2003; Christian et al., 1986). Here, cores and home range size varied with breeding phenology. The shift to smaller home ranges during late breeding season (Fig. 1 & 6) was consistent with our expectation of shifts in space use for floaters and breeders. During late breeding season, nestlings require less immediate attention (e.g., brooding or shading). Consequently, females are able to more actively defend their territories, potentially suppressing movements of floaters near territory boundaries, as well as more actively provision nestlings during a time when they require more food to support growth and development (Watson, 2010). The increase in number of core areas in late breeding season for territorial eagles is also consistent with added home range structure resulting from this change.

Within these different home range cores, however, we found that individuals selected habitats and resources differently (Fig. 2), which aligns with our predictions about individuals partitioning space for different activities (e.g., roosting and foraging).

We did not predict, however, the low among-individual variance in habitat and resource selection. While among-floater variance was slightly greater—likely due to their greater flexibility in movement strategy compared to territorial individuals—there was overall low individual heterogeneity (Fig. 5). This is somewhat in contrast to other work that highlights the importance of accounting for individual variation in resource selection studies (Lesmerises and St-Laurent, 2017) and recent findings of high heterogeneity in golden eagle selection for certain landscape features during migration (Eisaguirre et al., 2019b). Our findings herein suggest that it may be just as important to condition inference of habitat and resource selection patterns on the structure of individuals’ home ranges, as it is on the individuals. Further, the extremely low individual heterogeneity among territorial individuals suggests that the realized niche of territorial eagles during the breeding season in this area is relatively narrow. Thus, territory occupancy and, thereby, reproductive performance of the population is sensitive to the distribution of suitable habitat, perhaps in addition to dynamic energy subsidies, such as uplift.

### Implications & conclusions

Given the importance of floaters to the population ecology of many long-lived taxa, our work sheds light on the nuances of space use and partitioning of dynamic and static habitat resources between floaters and territorial individuals. The comparatively expansive movements of floaters and their proximity to territorial individuals—both in spatial and habitat distances—suggests the potential for some of the complex density-dependent effects an underworld of floaters can have on breeders, including competition for food, alteration of breeder behavior through territory intrusions, and reduced survival through fatal conflicts (Hunt, 1998; Ferrer et al., 2004; López-Sepulcre and Kokko, 2005; Carrete et al., 2006a; Bretagnolle et al., 2008; Watson, 2010; Ferrer et al., 2015). We may also conclude that habitat and landscape conservation plans that prioritize areas known to encompass golden eagle territories in this, and likely other, study areas also help conserve floaters and facilitate demographic buffering.

While the importance of floaters to the populations of long-lived, territorial species is understood (Penteriani et al., 2011), pre-breeding-age individuals, although not included in our sample, here, should also not be ignored; they must survive to become floaters and eventually breeders. Pre-breeding age individuals (i.e. typically those in their first to third summers for golden eagles) can have markedly different movement and space use patterns, as they are not yet attempting to enter the breeding population (Delgado and Penteriani, 2008; Delgado et al., 2009; Caro et al., 2011; McIntyre and Lewis, 2018). Once they enter the floater demographic, they, as we have shown, utilize areas interstitial to or briefly within territories, exhibiting different movements but only slight differential resource selection compared to territorial breeders. This transition, therefore, likely marks the onset of one to several breeding seasons of balancing time and effort between awaiting a territory vacancy, usurping a territorial resident, maintaining access to food, and learning about dynamic resources available within candidate breeding territories that would enhance future reproductive success.

## Acknowledgements

T. & D. Hawkins, M. Kohan, B. Robinson, and many others provided support in the field, and J. Liguori and N. Paprocki helped age eagles. C. McIntyre, K. Kielland, and P. Doak provided excellent feedback that helped improve this research and manuscript. To all of these friends, we are most grateful. Funding was provided by the Alaska Department of Fish & Game (ADF&G) through the federal State Wildlife Grant Program, and the U.S. Fish & Wildlife Service (USFWS) provided additional PTTs and data. JME was supported by the Calvin J. Lensink Fund during part of the project. The findings and conclusions of this paper are those of the authors and do not necessarily represent the views of the USFWS.

## Data accessibility

All movement data used for this manuscript are archived in the online repository Move-bank (https://www.movebank.org/; IDs 17680093 and 19389828). The data contain information considered confidential and sensitive by the State of Alaska (State Statute 16.05.815(d)), but they could be made available for research at the discretion of the Alaska Department of Fish & Game and U.S. Fish & Wildlife Service.

## Appendix 1

## Habitat types

**Table 2:** Habitat types used in analysis.

| AKVWC class | habitat type |
| --- | --- |
| Bareground | bare |
| Freshwater or Saltwater | water |
| Bareground (Beach or Tide Flat) (Southern Alaska),<br>Herbaceous (Marsh) (Interior Alaska, Cook Inlet Basin),<br>Herbaceous (Marsh) (Northern and Western Alaska),<br>Herbaceous (Tidal) (Southern Alaska), Herbaceous (Wet-Marsh) (Southern Alaska), Herbaceous (Aquatic), Low<br>Shrub (Tidal) (Southern Alaska), Herbaceous (Wet-Marsh)<br>(Tidal) | wet |
| Herbaceous (Mesic) (Interior Alaska, Cook Inlet Basin),<br>Herbaceous (Mesic) (Northern and Western Alaska), Herba-<br>ceous (Mesic) (Southern Alaska), Herbaceous (Peatland)<br>(Southern Alaska), Herbaceous (Wet) (Interior Alaska,<br>Cook Inlet Basin), Herbaceous (Wet) (Northern and West-<br>ern Alaska), Lichen, Moss, Moss (Southern Alaska), Sparse<br>Vegetation (Interior Alaska, Cook Inlet Basin), Sparse Veg-<br>etation (Northern and Western Alaska), Tussock Tundra<br>(Low shrub or Herbaceous), Fire Scar | open |
| Low Shrub, Low Shrub (Peatland) (Southern Alaska),<br>Dwarf Shrub, Dwarf Shrub (Southern Alaska), Dwarf<br>Shrub-Lichen, Dwarf Shrub, or Herbaceous (Mesic) (South-<br>ern Alaska), Low Shrub or Tall Shrub (Open-Closed), Low<br>Shrub/Lichen, Low-Tall Shrub (Southern Alaska), Tall<br>Shrub (Open-Closed) | shrub |
| Deciduous Forest (Open-Closed), Deciduous Forest<br>(Open-Closed) (Seasonally Flooded) (Southern Alaska),<br>Deciduous Forest (Woodland-Closed) (Southern Alaska),<br>Hemlock (Woodland-Closed), Hemlock-Sitka Spruce<br>(Woodland-Closed), Needleleaf Forest (Open-Closed) (Sea-<br>sonally Flooded) (Southern Alaska), Needleleaf Forest<br>(Woodland-Open) (Peatland) (Southern Alaska), Sitka<br>Spruce (Woodland-Closed), White Spruce or Black Spruce<br>(Open-Closed), White Spruce or Black Spruce (Woodland),<br>White Spruce or Black Spruce-Deciduous (Open-Closed),<br>White Spruce or Black Spruce/Lichen (Woodland-Open) | forest |
| Urban, Agriculture, Road | human |
| Ice-Snow | ice |

## Area within 95% contour of kernel density estimates

Median (IQR) for territorial eagles: 55 (37, 126) km^2^

Median (IQR) for floater eagles: 5483 (3110, 13974) km^2^

## Notes

### Competing Interest Statement

The authors have declared no competing interest.

